# Neutralization of European, South African, and United States SARS-CoV-2 mutants by a human antibody and antibody domains

**DOI:** 10.1101/2021.03.22.436481

**Authors:** Zehua Sun, Andrew Kim, Michele D Sobolewski, Nathan Enick, Chuan Chen, Cynthia Adams, Jana L Jacobs, Kevin D McCormick, John W Mellors, Dimiter S Dimitrov, Wei Li

**Author notes:** Corresponding authors: Zehua Sun,; Wei Li.

## Abstract

Severe Acute Respiratory Syndrome Coronavirus-2 (SARS-CoV-2) transmission with several emerging variants remain uncontrolled in many countries, indicating the pandemic remains severe. Recent studies showed reduction of neutralization against these emerging SARS-CoV-2 variants by vaccine-elicited antibodies. Among those emerging SARS-CoV-2 variants, a panel of amino acid mutations was characterized including those in the receptor-binding domain (RBD) of the SARS-CoV-2 spike (S) glycoprotein. In the present study, we evaluated our previously identified antibody and antibody domains for binding to these RBD variants with the emerging mutations, and neutralization of pseudo typed viruses carrying spike proteins with such mutations. Our results showed that one previously identified antibody domain, ab6, can bind 32 out of 35 RBD mutants tested in an ELISA assay. All three antibodies and antibody domains can neutralize pseudo typed B.1.1.7 (UK variant), but only the antibody domain ab6 can neutralize the pseudo typed virus with the triple mutation (K417N, E484K, N501Y). This domain and its improvements have potential for therapy of infections caused by SARS-CoV-2 mutants.

## Introduction

Antibodies targeting the SARS-CoV-2 spike protein receptor-binding domain (RBD) are developed as therapeutic for neutralization. Recently, the prevalent emerging SARS-CoV-2 variants have rapidly generated mutations to escape of neutralization of antibodies, both in non-ACE2 binding residues and RBD residues in the spike protein [1, 2]. The known emerging variants include those that were identified in the United Kingdom (lineage B.1.1.7), South Africa (501Y.V2 also known as B.1.351) and Brazil (P.1) [3]. The reported lineage B.1.1.7 variant contains 8 mutations which are located in S protein (Δ69-70, Y144Del, N501Y, A570D, P681H, T716I, S982A and D1118H), with one mutation N501Y located in RBD. The B.1.351 lineage variant which was rapidly identified as a predominant circulating genotype in South Africa was reported containing 9 mutations in the S protein (L18F, D80A, D215G, L242-244del, R246I, K417N, E484K, N501Y and A701V), three of which (K417N, E484K and N501Y) are in the RBD. The P.1 variant also has K417T, E484K, and N501Y mutations in the RBD. These mutations contribute to an increased affinity of the RBD for human angiotensin-converting enzyme 2 (ACE2), improving viral entry into cells [4].

The rapid emergence of new variants is of great concern for their increased transmissibility and evasion of neutralizing antibody therapeutics and the humoral response generated by currently approved vaccines including Pfizer BioNtech BNT162b2 lipid-nanoparticle-formulated, and the Moderna mRNA-1273 vaccine, which were both developed based on the earlier viral isolates [3, 5–7]. Reduced neutralization of SARS-CoV-2 lineage B.1.1.7 and other variants by convalescent and vaccine sera has been reported [8].

We previously reported a panel of potent neutralizing antibodies and antibody domains against the receptor-binding domain (RBD) of the SARS-CoV-2 spike (S) glycoprotein. An antibody domain ab6 specifically neutralized SARS-CoV-2 (wild type) with a 100% neutralization at 1.5 μg/ml as measured by two independent *in vitro* replication-competent virus neutralization assays [9]. Another antibody domain ab8 based high-affinity antibody potently neutralizes mouse-adapted SARS-CoV-2 in wild-type mice at a dose as low as 2 mg/kg [10]. One full length IgG1 antibody ab1 competes with human ACE2 for binding to RBD and potently neutralizes replication-competent SARS-CoV-2 in a replication-competent mouse ACE2-adapted SARS-CoV-2 in BALB/c mice and native virus in hACE2-expressing transgenic mice [11]. In the present study, we aim to examine the neutralization potency of these previously reported antibodies against the emerging variants containing key spike/RBD mutations by using comprehensive ELISA and pseudo typed virus neutralization assays. We found all of these three antibodies retains effective neutralization against the UK SARS-CoV-2 variants, with similar neutralizing IC50s as against the wild type virus. However, only ab6 neutralized SARS-CoV-2 pseudovirus carrying the SA triple RBD mutation. The results inform the further clinic development of these antibodies against emerging SARS-CoV-2 variants either by making antibody cocktails or by constructing bispecific antibodies.

## Results

Ab1, ab6, and ab8 are all antibody or antibody domains that can compete with human ACE2 for binding to the wild type SARS-CoV-2 RBD. In order to examine the binding affinity of these antibody and antibody domains to RBDs bearing different amino acid mutations, we engineered and purchased a panel of RBD variants bearing single amino acid mutation or combination of mutations (N354D/D364Y, and K417N/E484K/N501Y respectively) in each variant respectively, and performed a comprehensive ELISA assay (Figure 1).

**Figure 1.**
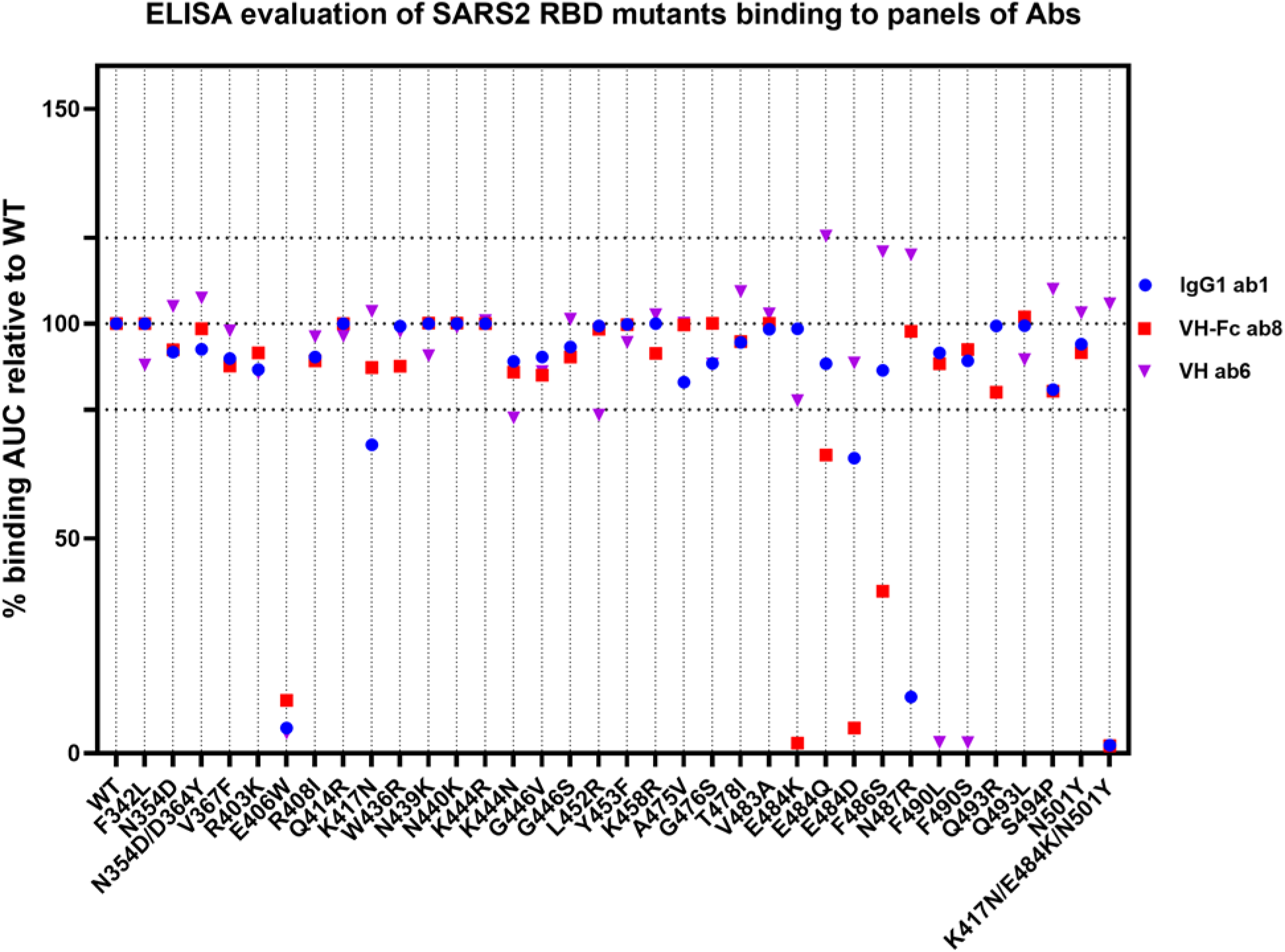
AUCs of three antibody and antibody domains to RBDs bearing different amino acid mutations, or combinations of specific mutations.

Results are showing that antibody VH domain ab6 can bind 30/33 single mutation variants including K417N, E484K and N501Y without any significant reduction of <u>AUC</u>. Worth of noted, the E484K substitution which is known to be present in the B.1.351 and P.1 variants has been reported as an escape mutation for a panel of characterized monoclonal antibodies including REGN10933 and Ly-CoV555 [12]. This E484K substitution also performed an escape for VH-Fc ab8, but not for IgG1 ab1. Consistently, E484Q and E484D substitutions [13] analog to E484K, can also fully or partially escape ab8. K417N, observed occurrence in variant dominantly transmitting in South Africa, is another key mutations in RBD that is reported to escape of neutralization from 33% mAbs tested in an assay, compared to wild typed SARS-CoV-2 (wild type). IgG1 ab1 showed decreased binding to this K417N mutation while ab8 and ab6 retain the binding. RBD variant bearing combination of K417N, E484K and N501Y can be recognized by VH ab6 without any reduction of binding AUC, but can completely block the binding of IgG1 ab1 and VH-fc ab8. These observations indicated that IgG1 ab1 and VH ab8 may completely loss the neutralization of reported B.1.351 lineage variant. However, both IgG1 ab1 and VH-Fc ab8 retain binding to the single mutation N501Y, indicating they may be still effective against the UK variants since N501Y mutations is the only mutation observed in RBD in UK strains. Naturally occurring mutation at a proximal position F490L and F490S [1] can completely block the binding of VH ab6, indicating F490 an important residue for ab6 binding. Another recent variant B.1.429 emerged in California, United States, contains a single L452R RBD mutation [14]. Our result showed L452R in RBD does not significantly block the binding of ab1, ab6 or ab8 (percent binding AUC relative to WT all above 75%). A single mutation, E406W [15], allowed for escape from REGN antibodies, also escaped from ab1, ab6 and ab8.

Next we used the S1 and full-length S trimer proteins carrying the whole sets of spike mutations (both RBD and non-RBD mutations) in UK, South Africa (SA) and Brazil variants to evaluate binding of these three antibodies by ELISA (Figure 2). SA and Brazil variants can completely deplete the binding of ab1 and ab8, but failed to escape from ab6.

**Figure 2.**
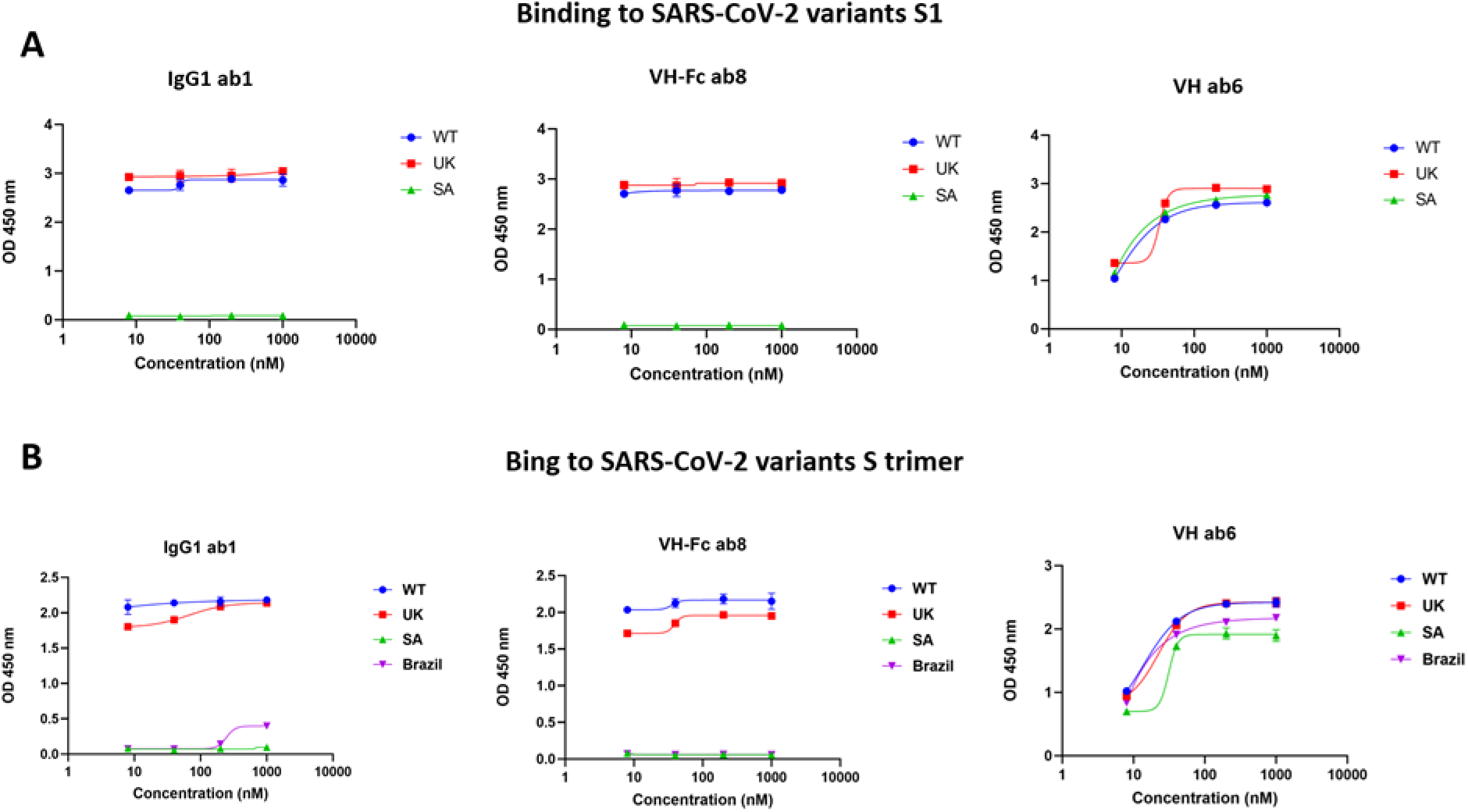
ELISA of ab1, ab6 and ab8 against S1 and full-length S trimer proteins carrying the whole sets of spike mutations (both RBD and non-RBD mutations) in UK, South Africa (SA) and Brazil respectively.

In order to validate the binding results are corroborating with neutralizing activities of our antibodies, we generated pseudo typed viruses carrying S proteins carrying mutations of B.1.1.7 lineage variant and the mutations in RBD of SA variant (K417N, E484K, and N501Y). Using HIV-1 lentiviral backbone pseudo-typing, we assessed the neutralization activity of these antibodies. Ab6 can neutralize both pseudo typed variants bearing different mutations, giving a 9.4 nM of IC50 in neutralizing pseudo typed B.1.1.7 lineage variant, and a 4.2 nM of IC50 in neutralizing the SA RBD triple-mutant variant. By contrast, ab1 and ab8 can only neutralize B.1.1.7 lineage variant without neutralization of SA RBD triple-mutant variant. The UK variants neutralization IC50s were ~ 6.2 and 5.5 nM, respectively for ab1 and ab8, which is visibly less than the IC50s against wild type SARS-CoV-2 (0.5-2 nM for ab1, 0.5-3 nM for ab8) but still effective in therapeutic-accessible concentrations (Figure 3).

**Figure 3.**
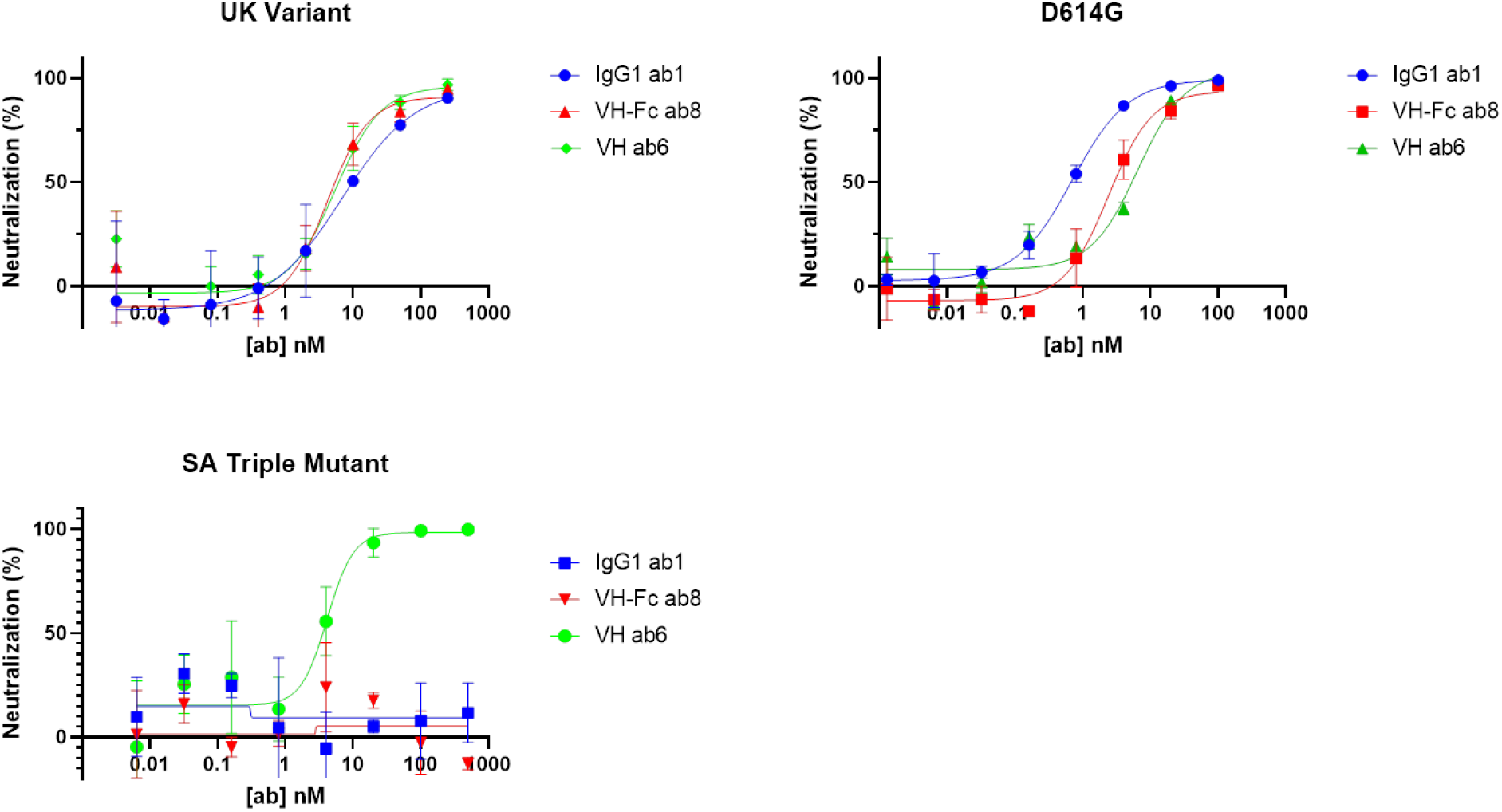
Neutralization assay of our antibodies against pseudo typed viruses carrying S proteins carrying different mutations.

## Discussion

Blocking the binding of viral receptor-binding domain (RBD) to human angiotensin-converting enzyme 2 (ACE2) receptor is one major solution of neutralization. The emerging SARS-CoV-2 virus variants bearing mutations in RBD decreased the efficiency of neutralization of FDA approved antibody therapeutics and vaccination induced humoral immune response [12]. Aiming to evaluate the efficacy of our previously reported antibodies in binding and neutralizing emerging mutant variants, we performed both ELISA assays against a panel of RBDs bearing different mutations and neutralization assays by using lentiviral virions pseudo typed by the variant spike proteins. Though it is possible that mutations outside the spike protein can determine or enhance escape from antibody neutralization, in most cases pseudotyped virus assays can reflect the result of a live replication competent system.

Recent studies show antibodies elicited by the Moderna 1273 mRNA vaccine decrease titers by 2.7-fold for B.1.1.7 variant and 6.4-fold for the viruses containing the B.1.351 spike protein [16, 17]. B.1.351 variant was observed markedly resistant to neutralization by convalescent plasma and vaccinee sera [18]. In a study of antibody response against the B.1.1.7 variant after a second dose of BNT162b2 vaccine, a modest reduction was observed in neutralization against pseudo typed viruses containing B.1.1.7 Spike mutations, and a significant additional loss of neutralization was observed against B.1.1.7 viruses bearing the Spike E484K mutation by BNT162b2 mRNA-elicited antibodies, convalescent sera and mAbs [19].

N501Y has little effect on neutralization by vaccinee sera as reported [20], however, 35% RDB-specific mAbs isolated from mRNA-1273 vaccinees showed a 100-fold or even higher loss of neutralization against N501Y mutant [18]. E484K mutant resulted a 3-6 fold reduction in neutralization, and 50% of the RBM specific mAbs lost neutralization against E484K mutant in one assay. K417N is another key mutations in RBD that can escape of neutralization from 33% mAbs tested in an assay. E484K in combination of N501Y and K417N can cause an additional loss of neutralization observed in convalescent sera [21, 22]. Although some of these variants and point mutations cause only partial reduction in neutralization by vaccine sera or currently approved mAbs, there is a risk that the constantly evolving variants could become dominant and evade current therapeutics and vaccines with higher efficiency. Antibody domain ab6 exhibited potency in neutralizing all these pseudo typed variants bearing these mutations. Interestingly, emerging variants including the 501Y.V2 and B.1.1.7 lineages also accumulated mutations and deletions in N-terminal domain (NTD) [23], and NTD-specific antibodies is comparable less resistant to mutant variants than RBD-specific antibodies.

Vaccines remain an efficient strategy for a long term control of the transmission of SARS-CoV-2 variants. Production and distribution of vaccines around the world require time and large scale coordination of logistics, allowing the virus to continue to spread and harbor new mutations. Our study provides highly potent antibody candidates that neutralized emerging mutant variants that can be used as emergency therapeutics where current vaccines are not present or less effective in prevention.

## Methods

### ELISA Assay

Ninety-six-well plates were coated with the RBD mutants (Sino Biological) at a concentration of 4ug/ml (diluted with 1xPBS) and incubated at 4C overnight (50ul per well). The next day, blocking was conducted with 100ul per well of 3% milk solution made from Bio-Rad nonfat dry milk powder and 1xPBS. The plate was incubated at room temperature for one hour. Primary antibodies were diluted with 3% milk blocking buffer and 1:5 serial dilutions starting at 1nm. After addition of 50ul of primary antibody dilutions to each well, plates were incubated at room temperature for 2 hours. The plate was subsequently washed with 0.05% Tween 1xPBS (PBST) solution using a plate washer (200ul per well, 4 rounds of washing). The secondary antibody was prepared with the 3% milk at a dilution of 1:1000. The secondary antibody was added (50ul per well) and incubates at room temperature for 1 hour. After 1 hour, the plate was washed as before and 50ul per well of TMB substrate (Sigma) was added. After 1-2 minutes, color development was stopped with 1M H2SO4 (50ul per well) and the plate was read at 450nm, which was plotted against antibody concentration and fitted by the non-linear regression curves fitting ([Agonist] vs. response---variable slope-four parameters), and the Area Under Curve (AUC) were calculated using GraphPad Prism 9.0.2.

### SARS-CoV-2 Pseudovirus Neutralization Assay

PSV was generated in 293T cells by co-transfection of pFC37K-CMV-S, an enhanced expression plasmid encoding for codon-optimized full-length SARS-CoV-2 S with the N-term HiBit tag removed, and pNL4-3.luc.R-E-mCherry-luciferase, an envelope deficient HIV-1 dual reporter construct that was cloned by recombination of the pNL.luc.R-E-plasmid (NIH AIDS Reagent Program) and the fully infectious pNL4-3 mCherry luciferase plasmid (Addgene) [24, 25]. After harvest, PSV was centrifuged at 500xg for 10 mins and supernatant removed and filtered with a 0.45 um syringe filter to remove producer cells. For neutralization assays 104 293T-hACE2 cells were plated in 100uL media per well in 96 well white-wall white-bottom plates (Perkin Elmer) and incubated overnight at 37°C. Ab6 was serially diluted 5-fold and incubated with 50uL of PSV for 1 hour at 37°C. After incubation, media was removed from wells containing 293T-hACE2 cells and replaced with PSV/mAb and spinoculation was performed at 1,000xg for 1 hour at RT. Plates were then incubated for 48 hours at 37°C. After 48 hours, plates were analyzed for luciferase production by adding 100uL of BriteLite Plus reagent (Perkin Elmer), incubating at RT for 2 minutes, and reading on a Victor Nivo microplate luminometer (Perkin Elmer).

## Author contributions

Z.S., D.S.D. and W.L. designed research; A.K., M.D.S., N.E., J.L.J and K.D.M performed research; C.C. contributed wild typed RBD protein; Z.S. isolated and characterized ab6; W.L. isolated and characterized ab1 and ab8; Z.S., J.W.M., D.S.D. and W.L. analyzed data; and Z.S., C.A., W.L., and D.S.D. wrote the paper.

## Acknowledgement

We thank the members of the Center for Antibody Therapeutics: Dontcho V Jelev, Du-San Baek, Xianglei Liu, Xiaojie Chu, Yae-Jin Kim and Megan Shi for their helpful discussions. This work was supported by the University of Pittsburgh Medical Center.

